# Extended kinship analysis of historical remains using SNP capture

**DOI:** 10.1101/2020.09.17.300715

**Authors:** Erin M. Gorden, Ellen M. Greytak, Kimberly Sturk-Andreaggi, Janet Cady, Timothy P. McMahon, Steven Armentrout, Charla Marshall

## Abstract

DNA-assisted identification of historical remains requires the genetic analysis of highly degraded DNA, along with a comparison to DNA from known relatives. This can be achieved by targeting single nucleotide polymorphisms (SNPs) using a hybridization capture and next-generation sequencing approach suitable for degraded skeletal samples. In the present study, two SNP capture panels were designed to target ~25,000 (25K) and ~95,000 (95K) nuclear SNPs, respectively, to enable distant kinship estimation (up to 4^th^ degree relatives). Low-coverage SNP data were successfully recovered from 14 skeletal elements 75 years postmortem, with captured DNA having mean insert sizes ranging from 32-170 bp across the 14 samples. SNP comparison with DNA from known family references was performed in the Parabon Fx Forensic Analysis Platform, which utilizes a likelihood approach for kinship prediction that was optimized for low-coverage sequencing data with cytosine deamination. The 25K panel produced 15,000 SNPs on average, which allowed for accurate kinship prediction in 17 of the 21 pairwise comparisons. The 95K panel increased the average SNPs to 42,000 and resulted in two additional pairwise comparisons (19 of 21). This study provides the groundwork for the expansion of research involving compromised samples to include SNP hybridization capture.

## 1. Introduction

Forensic laboratories are beginning to explore the use of large-scale single nucleotide polymorphism (SNP) panels for human identification in routine casework. The interrogation of a single-base target increases the likelihood of successful genotyping for degraded DNA samples, which is particularly beneficial for cases involving missing persons and historical remains with poor quality DNA. SNPs are known to have a low mutation rate making them valuable for distant kinship analysis [1], empowering kinship comparisons amongst distant relatives and facilitating the expansion of suitable references beyond immediate family members. This is of particular importance in decades-old cases, where generational gaps caused by deficient pedigrees and the passage of time may prevent closely-related references from being obtained.

Recent attention on SNPs in forensics has been primarily driven by the use of next-generation sequencing (NGS), which provides a means of generating massive amounts of data at one time for a large number of SNP loci. Commercially available NGS forensic kits, such as the ForenSeq DNA Signature Prep Kit (Verogen, San Diego, CA) and the Applied BioSystems Precision ID panels (Thermo Fisher Scientific, Waltham, MA), incorporate 100-200 SNPs for identity, phenotype and biogeographical ancestry inferences. Expanded PCR-based panels have been released for distant kinship estimation. These include Verogen’s ForenSeq Kintelligence panel of ~10,000 markers as well as an identity panel of ~1200 tri-allelic SNPs utilized with the QIAseq method (QIAGEN, Hilden, Germany) [2,3]. Yet, the nature of DNA that can be recovered from severely compromised samples (e.g., fragmented, damaged, and low endogenous content) limits the applicability of these amplicon-based tools. And furthermore, the bioinformatic pipelines used in conjunction with amplicon data require sufficient read depths to directly call genotypes at each SNP and thus may not be suitable for routine casework involving severely compromised samples.

Alternatively, high-density genotyping arrays targeting up to a million or more SNPs have been used recently in cold cases to obtain genome-wide data from unknown crime scene samples for investigative genetic genealogy. Upload of these forensic profiles to publicly available genealogy DNA databases, such as GEDmatch [4] and FamilyTreeDNA [5], has facilitated new leads and identified suspects where other lines of investigation have been exhausted (e.g., [6–8]). SNP arrays have therefore provided the key to unlock genetic genealogy as a forensic investigative tool. However, while the SNP arrays utilized for genetic genealogy by direct-to-consumer (DTC) DNA testing companies have been used on a wide variety of forensic case samples, including bone [9], they require high quantities of intact human DNA (≥ 1 ng) and are hindered by severe degradation and microbial contamination. Thus SNP arrays are not effective for highly degraded and/or environmentally challenged samples. For this reason, scientists are pursuing whole genome sequencing (WGS) as an alternative means to collect genome-wide SNP data from skeletal remains and other challenging samples. In one recently published study, WGS was utilized to generate SNP genotypes from unidentified human remains, providing information to help determine the identity of the unknown victim [10]. The drawback to WGS is that it is expensive, especially when sequencing samples with a high percentage of microbial DNA that is co-extracted from the human bone, such as in historical cases and other unidentified skeletal remains [11]. To maximize the usage of WGS data, scientists are now applying imputation methods [12–14] to produce complete genotypes from low coverage genomes in forensic cases [15].

Hybridization capture targeting the mitochondrial genome and small (~1200) SNP sets for identity testing has been successfully applied to forensic casework, including degraded and chemically treated samples [16–19]. Yet due to the time and cost involved in hybridization capture, this method is not widely used in forensic casework, which can largely be solved using routine short tandem repeat (STR) typing for human identification/kinship analyses or SNP arrays for genetic genealogy. Therefore, hybridization capture is, at the time of writing, most often reserved for the poorest quality samples or for those cases involving historical remains. Much broader use of hybridization capture is seen within the field of ancient DNA (aDNA). Such archaeogenomic research has shown the ability to obtain SNP genotypes from aDNA samples using hybridization capture and demonstrated that such data can be used to establish relationships between individuals and groups of individuals [20–22]. However, the methods involved in aDNA studies are not easily implemented within regulated forensic laboratories. For example, the SNP probes may be made in-house and are not commercially available, the software used to perform data analysis is not available to external users, or the amount of sequencing required is too high for routine casework. Consequently, the gains made within the field of aDNA to advance SNP capture from skeletal remains are not directly transferrable to forensic practitioners.

The goal of this study was to obtain thousands of nuclear SNPs from previously identified historical remains for kinship comparison to known references. Familial relationships ranging from parent/offspring to 4^th^ degree relatives were represented in the 14 cases. The generation of SNP profiles used for this study was achieved via three methods: WGS and two levels of targeted nuclear SNP capture panels (~25,000 SNPs and ~95,000 SNPs). Kinship analysis was completed using the Parabon Fx Forensic Analysis Platform (Parabon NanoLabs; Reston, Virginia) [23]. An evaluation of the efficacy of the three SNP approaches to achieve strong kinship support was performed, which would ultimately aid in the identification of unidentified human remains.

## 2. Materials and Methods

### 2.1 Non-probative casework samples

#### 2.1.1 Sample selection

Fourteen previously identified, non-probative skeletal samples from the Armed Forces Medical Examiner System’s Armed Forces DNA Identification Laboratory (AFMES-AFDIL) were selected for use in this study (Table 1). These specimens originated from a variety of case contexts, and all were approximately 75 years postmortem. Samples were selected based on the availability of family reference samples (FRS) for kinship comparison.

**Table 1.**
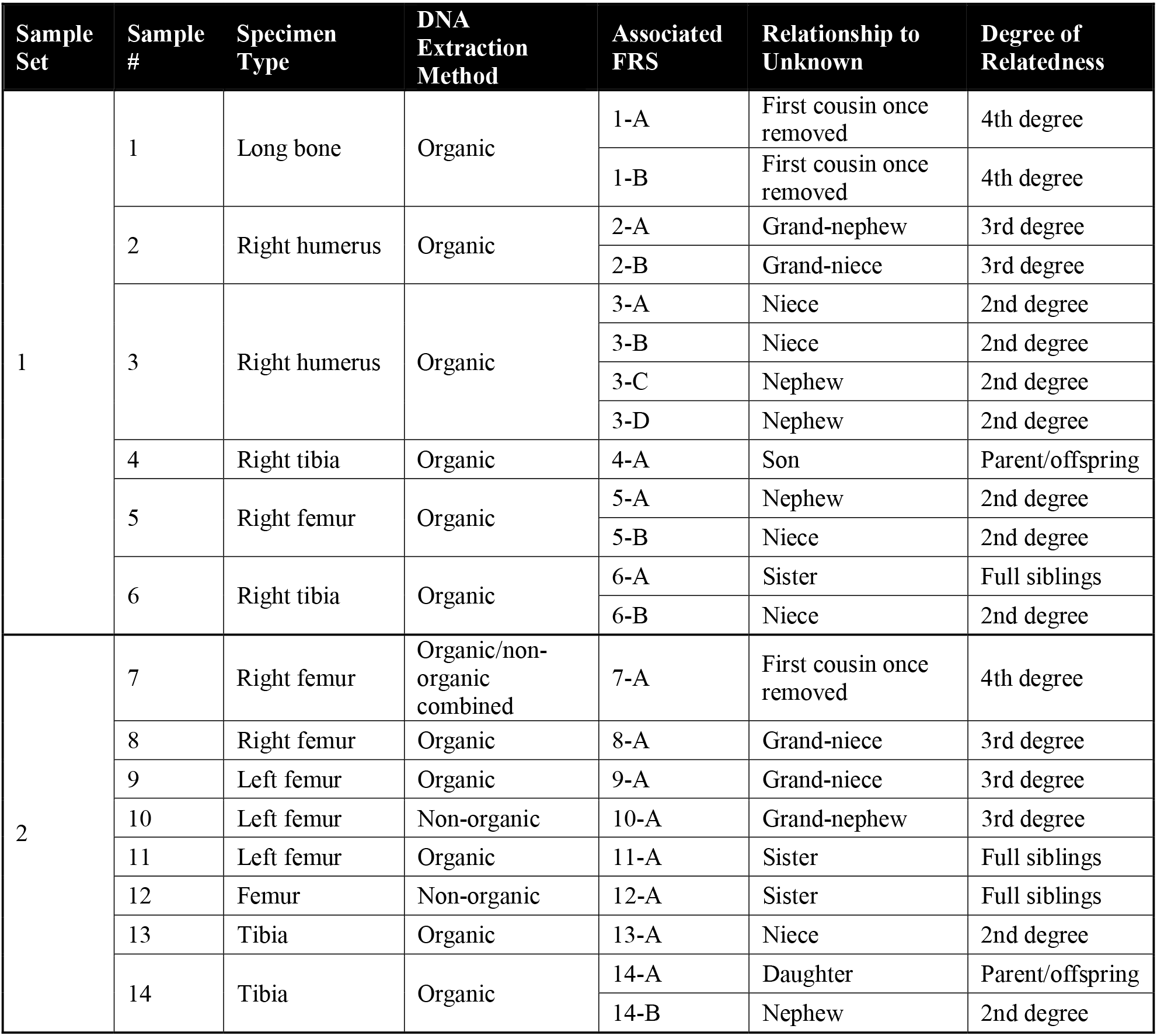
Sample information and laboratory processing procedures for 14 non-probative casework samples used in this study. Relationship information for 22 family reference samples (FRS) is also listed. Non-probative samples are divided into two sample sets for laboratory processing and sequencing. An example pedigree demonstrating familial relationships is presented in Figure S1.

#### 2.1.2 DNA extraction and repair

DNA was extracted from fragments of bone ranging from 0.2-1.0 g. Samples were first powdered and demineralized overnight at 56°C in a buffer containing 0.5 M EDTA, 1% lauroylsarcosine and 100 μl of 20 mg/ml proteinase K. DNA purification was achieved using either an organic protocol or the QIAquick PCR Purification Kit (QIAGEN) as described in [24]. One extraction control (reagent blank; RB) was included in each set (RB1 for set 1 and RB2 for set 2). In one instance (sample 7), previously generated DNA extracts were combined prior to DNA repair due to low volumes in order to maximize the DNA input for downstream processing.

A DNA repair step was performed on all DNA extracts, including RBs, using the NEBNext FFPE DNA Repair Mix (New England BioLabs Inc, NEB; Ipswich MA). Purification of the repaired DNA extract was performed using the QIAGEN MinElute PCR Purification Kit, followed by elution in sterile Tris-EDTA [10 mM Tris, (pH 7.5) 0.1 mM EDTA]. DNA was quantified with the dsDNA High Sensitivity (HS) Assay Kit on the Qubit 2.0 or Qubit 3.0 Fluorometer (Thermo Fisher Scientific) to determine input into NGS library preparation.

#### 2.1.3 Library preparation

The KAPA Hyper Prep Kit (Roche Sequencing, Pleasanton, CA) was used for library preparation. To evaluate the success of the library and capture procedures, a positive control (PC) was initiated at the beginning of library preparation for each sample set (PC1 for set 1 and PC2 for set 2). Adapter ligation was completed using dual-indexed adapters (Integrated DNA Technologies, Coralville, IA) matching sequences used in the Illumina TruSeq HT kits at an adapter concentration of 15 μM. Ten cycles of PCR amplification were completed on a GeneAmp PCR System 9700 thermal cycler (Applied Biosystems, Foster City, CA) following the manufacturer’s recommendations. Samples were eluted in 25 μl of Tris-EDTA following bead-based cleanup. Libraries from all sample sets were quantified on the 2100 Bioanalyzer instrument with the Agilent DNA 7500 Kit (both Agilent Technologies, Santa Clara, CA).

#### 2.1.4 25K and 95K SNP capture panel design

Two custom SNP panels were designed to allow overlap with the Infinium CytoSNP-850K v1.1 BeadChip (Illumina, San Diego, CA) for extended kinship comparisons. The first panel contained 24,999 genomic SNPs (Table S1; referred to as the 25K SNP panel) and the second panel included 94,752 genomic SNPs (Table S2; referred to as the 95K SNP panel). Criteria for SNP target selection included likelihood to perform well for kinship (i.e. high minor allele frequency in all populations, low F_ST_, and low linkage disequilibrium between SNPs), distribution across the genome, and the GC content of the surrounding genomic region (25-60%). A majority of the SNPs chosen were bi-allelic, and while not specifically targeted for their polymorphic status, smaller proportions were tri- and tetra-allelic (Table S3). The presence of candidate SNPs in commercially available microarray kits ensured kinship comparison could be performed with samples genotyped using microarray platforms (i.e. high quality FRS).

#### 2.1.5 Hybridization capture of SNP panel targets

Hybridization capture of the 25K and 95K SNP panel targets was facilitated by myBaits Custom DNA-Seq kits (Arbor Biosciences, Ann Arbor, MI). Bait design for the 25K panel consisted of four baits per SNP (~100,000 unique RNA baits) and the 95K SNP panel was designed with one or two baits per SNP (~180,000 unique RNA baits). The hybridization conditions followed the manufacturer’s recommended protocol [25] with a 24-hour hybridization at 65°C using a Veriti thermal cycler (Thermo Fisher Scientific). An attempt was made to target the maximum recommended DNA input into capture (500 ng). For some samples, capture input was reduced for the 95K panel due to lack of remaining library (Table S4). The captured product was split into two portions and both portions were amplified independently for 19 PCR cycles with KAPA HiFi HotStart ReadyMix (Roche Sequencing). The two amplified capture products for each sample were combined and purified with the MinElute PCR Purification Kit, eluting in 25 μl of Tris-EDTA. Purified capture product was quantified using the 2100 Bioanalyzer instrument with the Agilent DNA 7500 Kit.

#### 2.1.6 Library pooling and sequencing

Each sample set was sequenced separately by pooling an equal volume of captured libraries within the set (excluding the PC). The PCs were not sequenced, as previous studies have shown sequencing of PCs with low quality samples can cause instrument crosstalk and complicate data interpretation [17,26,27]. Pool molarity was determined using the Agilent DNA 7500 Kit on the 2100 Bioanalyzer instrument. Pools were loaded for sequencing at 10 pM and spiked with PhiX Sequencing Control v3 (Illumina) at a 2.5% concentration. Sequencing was performed on a MiSeq FGx Forensic Genomic System (Verogen) in Research Use Only mode. Paired-end sequencing was completed using 300-cycle MiSeq Reagent Kits v2 (Illumina; 2 × 150 cycles).

To evaluate the performance of whole genome sequencing (WGS) as an enrichment-free SNP profiling option for better-quality samples as well as deeper sequencing of the 95K capture product, two sample libraries (2 and 5) were sequenced using a higher-throughput Illumina NextSeq 550. These libraries were individually normalized and pooled in equimolar concentration for each run. The pools were loaded at 1.45 pM (80% of the manufacturer’s 1.8 pM recommended loading concentration) with 1% PhiX control spiked-in as a sequencing control. Paired-end sequencing was completed using NextSeq 500/550 300-cycle High-Output Kits v 2.5 (Illumina; 2 × 150 cycles). High-Output kit used for the generation of the NextSeq data allows for over 25 times more sequence data to be generated compared to MiSeq v2 sequencing (800M versus 30M paired reads, respectively).

#### 2.1.7. Microarray genotyping

DNA extracts from two additional samples were sent to AKESOgen (Peachtree Corners, GA), a CLIA-certified sequencing laboratory, for microarray genotyping with the Illumina Infinium CytoSNP-850K v1.1 BeadChip. Both unrepaired and repaired DNA extracts were tested using two genotyping protocols. Genotyping was halted after the first four samples failed to generate any usable data (genotyping call rate <50%). A summary of these results is available in Table S5, but no further discussion will be provided below due to the extremely poor quality of the genotyping data.

### 2.2 Family reference samples

#### 2.2.1 Sample selection and DNA extraction

A total of 22 FRS were selected for use in this study based on known relationships to the previously identified case samples (Table 1). An additional 17 FRS with no relation to the case samples were also genotyped for a total of 39 FRS for comparison. All FRS donors provided informed consent for samples to be used in research and quality improvement activities. The use of these samples was approved by the Defense Health Agency Office of Research Protections (Protocol # DHQ-20-2073). FRS were saliva samples deposited on either Bode buccal collectors or cotton buccal swabs. The QIAamp DNA Investigator Kit (QIAGEN) was employed for DNA extraction following one of two manufacturer protocols: isolation of total DNA from FTA and Guthrie cards using two punches from Bode buccal collectors, or isolation of total DNA from surface and buccal swabs using one buccal swab. The final elution volume for all samples was 100 μl of Tris-EDTA. DNA was quantified using both the Qubit dsDNA HS Assay Kit on the Qubit 3.0 Fluorometer and the Plexor HY DNA Quantification Kit (Promega Corporation, Madison, WI) prior to submission for microarray genotyping.

#### 2.2.2 Microarray genotyping

FRS were genotyped at AKESOgen using the Infinium CytoSNP-850K v1.1 BeadChip. Genotypes were called using the GenomeStudio Software (Illumina) and the data were reviewed prior to upload to the Parabon Fx Forensic Analysis Platform for analysis.

### 2.3 Data analysis

#### 2.3.1 Non-probative case samples

Demultiplexed FASTQ files were generated from the raw data by the MiSeq Reporter software (Illumina) and imported into the Parabon Fx Forensic Analysis Platform. Reads were aligned to the human reference genome (GRCh38), and duplicate mapped reads were removed. Alignment parameters included penalties for mismatches (4), open gaps (6), extension gaps (1) and unpaired reads (9). Minimum map and base qualities were not required (i.e. set to 0) in order to allow for the maximum number of called SNPs, which are assessed using a probabilistic approach. The software implements a genotype likelihood approach for each locus under investigation, which facilitates the use of low coverage SNPs (≥ 1X) and maximizes the use of available information [28]. In short, the likelihood of the data under the hypothesis of each possible genotype is calculated given factors such as the number of reads per allele and the sequencing quality score. This approach increases the probative power of the analysis and avoids potentially erroneous genotype calls generated using an absolute calling threshold. To account for the effects of DNA damage typical of ancient and historical remains, cytosine deamination was assessed at the time of alignment [29,30]. This assessment is performed utilizing a different mapping (than the original alignment described above) as the read pairs are merged and aligned to the GRCh38 as single-end reads. This allows for estimation of the mismatch rate at the ends of the fragments, specifically cytosine to thymine error on the 5’ end and guanine to adenine error on the 3’ end that are indicative of cytosine deamination [29,30]. Based upon this assessment (e.g., Figure S2), the user has the option to correct for the observed damage. In addition to the use of the merged-pair mapping, the position within the fragment is incorporated into the likelihood calculations when generating the SNP profile when the damage correction option is selected. Due to the age of the specimens tested, damage correction was applied to all bone sample profiles generated herein. SNP genotype likelihood profiles were generated for each captured SNP set using read depth thresholds of 1X, 5X and 10X. Though no enrichment was performed, WGS data were analyzed for the 843,223 SNPs present on the Infinium CytoSNP-850K BeadChip at the 1X, 5X and 10X thresholds.

Ancestry inference was performed in Fx for both the unknown and reference samples, which was then used along with self-reported race (for the FRS donors) to determine the appropriate population allele frequency file for kinship calculations. Allele frequencies from the five populations represented in the 1000 Genomes Project [31] were used for the ancestry prediction. The Parabon Fx Forensic Analysis Platform estimates pairwise relatedness using the likelihood formulas derived in [32], adapted for the scenario of testing a specific set of kinship hypotheses by comparing genotype likelihoods to called genotypes from a reference sample. This approach takes into account all possible genotypes for the captured sample, as well as the genotype frequencies of the selected population. Kinship predictions were performed against the entire FRS dataset (Figure S3), and log likelihoods were calculated within the software for each degree of relatedness (DOR) up to 4^th^ degree (from self/monozygotic twin, parent-offspring, full sibling, 2^nd^ degree, 3^rd^ degree, and 4^th^ degree) as well as unrelated. Likelihood ratios (LRs) were generated within the software for the most likely DOR compared to the unrelated category. LRs over 10,000 were considered strong evidence of relatedness, based on previously established forensic guidelines [33].

#### 2.3.2 Family reference samples

FRS SNP genotypes were evaluated for genotyping call rate (i.e. the proportion of SNPs with called genotypes), and those below 70% were excluded from further analysis. The remaining FRS SNP genotypes were loaded into the Parabon Fx software, and each sample’s ancestry proportions were estimated. Due to the large number of SNPs recovered in the FRS microarray data, ancestry prediction was possible for seven well-defined global population groups [34]. In addition to kinship comparisons with the case samples, kinship between each FRS pair was predicted and compared to the expected relationships.

## 3. Results and Discussion

### 3.1 Non-probative casework sample assessment

A total of 16 libraries were prepared and sequenced: 14 bone samples and two RBs. The two PCs were not sequenced in order to minimize crosstalk but they were assessed during several quantification steps for library product as a quality control assessment (Table S4). Sequence data for both SNP capture panels (i.e. 25K and 95K) were generated on the MiSeq from each of the 16 libraries. Additionally, WGS data were produced from the two better quality libraries on the NextSeq (along with additional 95K capture data for the same samples). Metrics for each of the seven sequencing runs can be found in Table S6, and alignment and mapping metrics for all samples are presented in Table S7. Average DNA fragment length after capture was ~120 bp in the 25K and 95K data (Figure 1). The fragment length distributions varied by sample, but each showed a peak between 20-25 bp (Figure 2). These short reads break with the unimodal distribution of endogenous DNA fragments and are likely due to non-specific mapping of off-target capture product. Typically, reads <30 bp are excluded from sequence alignments in aDNA research (e.g., [35]), which minimizes the proportion of off-target read mapping. Such short read filtering may also be required for forensic casework involving degraded DNA, and this parameter can be incorporated into future versions of the Parabon Fx software. The maximum DNA fragment length was variable, as sample 3 (Figure 2a) had a maximum fragment length exceeding 400 bp, while sample 11 (Figure 2c) contained endogenous human DNA that was altogether <100 bp in length. Despite the skewed fragment length distribution due to the non-specific mapping of off-target short reads, these samples were shown to exhibit considerable DNA degradation consistent with expectations from historical remains.

**Figure 1.**
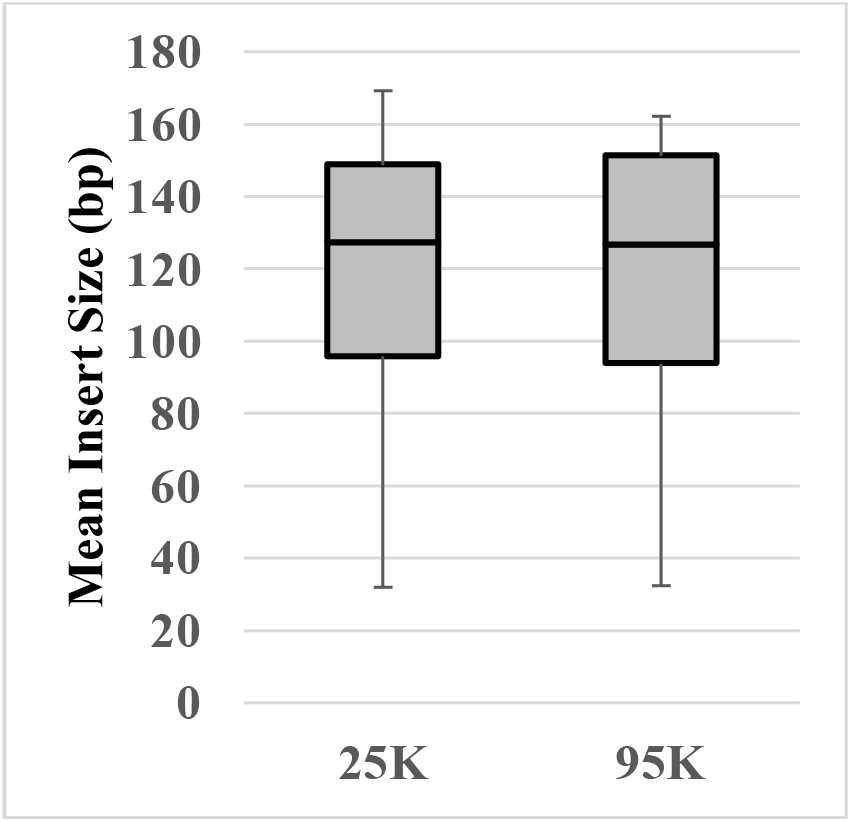
Distribution of the mean insert size (average DNA fragment length) in base pairs (bp) of reads mapped to the human genome (GRCh38) for the 25K and 95K SNP capture approaches.

**Figure 2.**
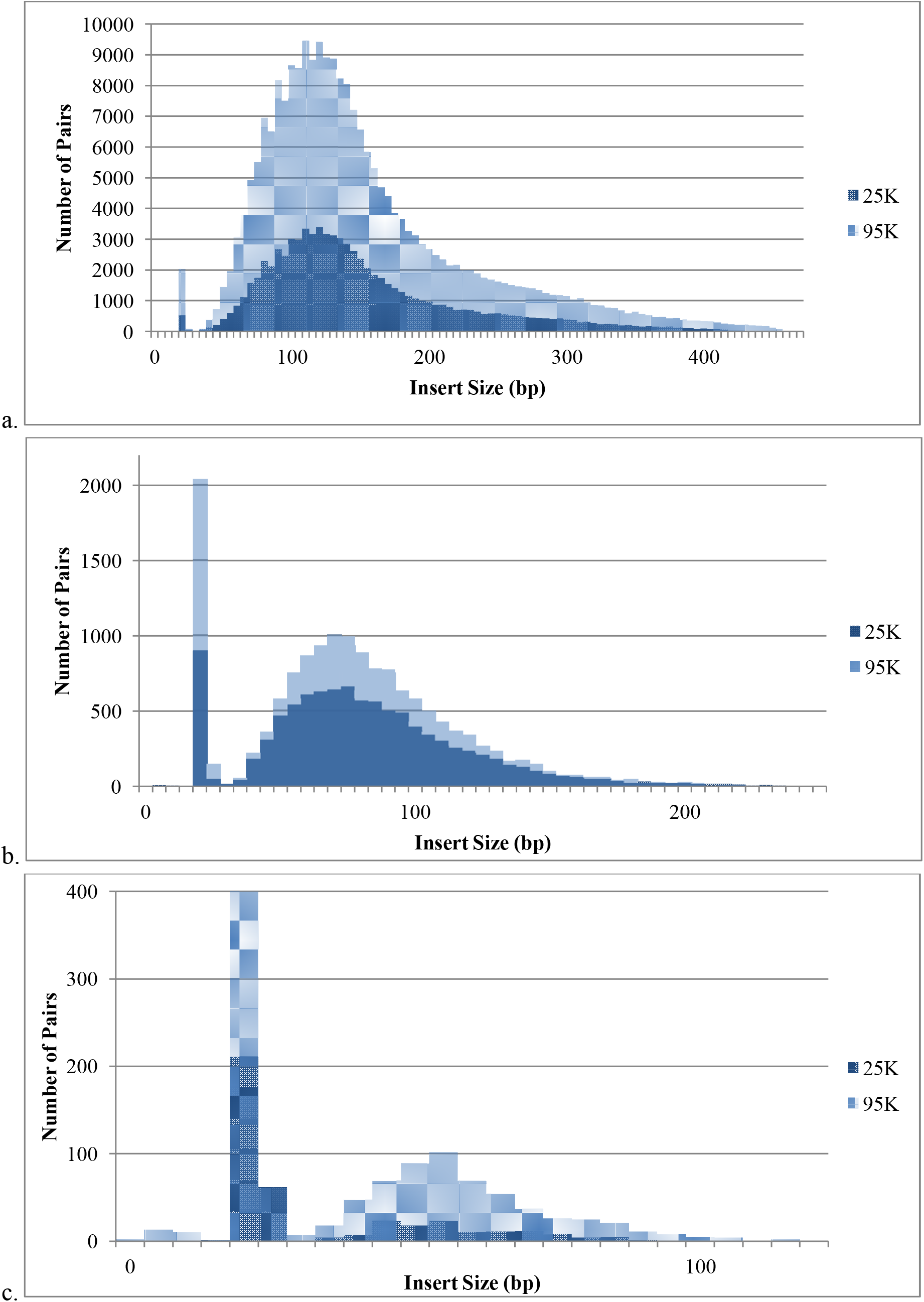
Fragment length (insert size) distribution for three exemplary samples using 25K and 95K capture. Samples represent the range of quality observed in this study: a) sample 3, b) sample 1, and c) sample 11. Note the difference in scale between samples for both the x and y axes. Peaks at 20-25 bp are likely due to non-specific mapping of off-target reads.

The proportion of unique reads that mapped to GRCh38 averaged ~45% for both 25K and 95K capture approaches (Table S7), ranging from ~30% to ~70% (Figure 3). In other words, even after capture to enrich for human SNPs, many of the historical bone samples tested here contained considerable environmental DNA in the sequenced libraries. To increase the endogenous human DNA proportion, a second round of capture could be performed. However, this adds time and expense to the procedure and should be considered on a case-by-case basis. The generally poor quality of the samples tested in this study is typical of specimens submitted to the AFMES-AFDIL for DNA testing [11,17,36], and likely explains the failed microarray testing discussed in 2.1.7. Though other, higher quality forensic casework samples may be amenable to microarray testing [9], severely compromised samples such as those from decades-old missing persons cases will likely necessitate alternative genotyping methods for SNP recovery such as hybridization capture.

**Figure 3.**
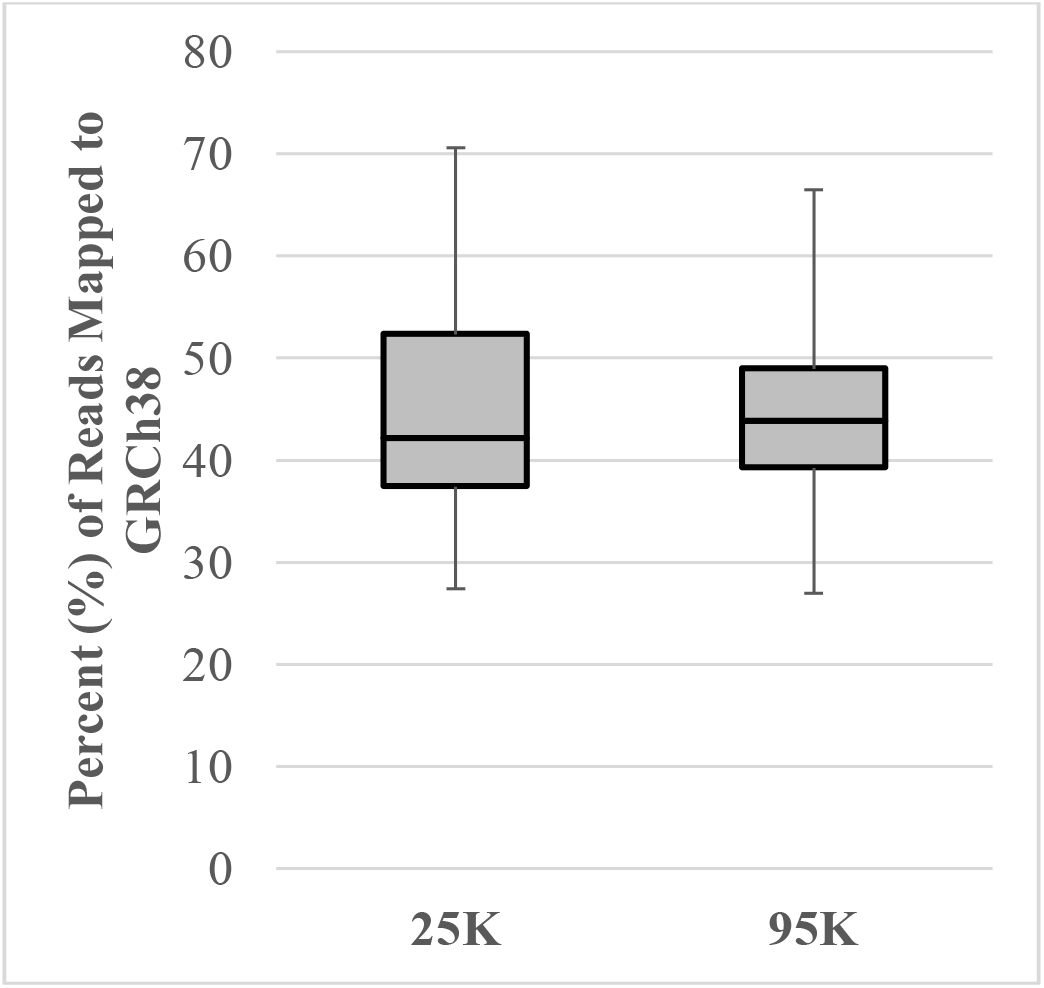
Distribution of the percent of unique reads mapped to the human genome (GRCh38) after 25K SNP capture and 95K SNP capture.

The MiSeq data resulted in an average of over 15,000 and 42,000 SNPs in the 25K and 95K panels, respectively. Sample 11 consistently performed poorly after capture using both panels, generating 159 SNPs from the 25K panel and 625 SNPs from the 95K panel (Figure 4a). Presumably, the relative failure of sample 11 is a consequence of the high degree of DNA degradation present as the mean insert size was only 32 bp, the smallest of all the samples tested (Figure 2c; Table S7). When comparing the performance of the 25K and the 95K panels, the raw number of SNPs recovered was greater using the 95K panel in 13 of 14 samples (92.9%). Moreover, ten of the 14 samples (71.4%) produced more than 25,000 SNPs when the 95K assay was used for capture (Figure 4a), which exceeds the maximum number of SNPs targeted by the smaller 25K panel. One sample (sample 4) produced fewer SNPs using the 95K capture assay when compared to the 25K capture assay; however, this can be attributed to the reduced DNA input for the 95K capture reaction due to insufficient remaining library volume (Table S4), rather than panel performance. It is expected that had the maximum library input into capture been attained, increased SNP recovery would have been observed. Interestingly, the proportional SNP recovery at 1X for each sample was similar between the 25K and 95K panels when the capture DNA input was equal (Figure 4b). The lowest SNP recovery (both in terms of absolute numbers but also proportion) was observed in samples 1, 4, 7 and 11. Three of these four specimens (4, 7 and 11) were subjected to post-mortem chemical treatment (formalin preservation), as described in [17]; thus, the low SNP recovery from these three chemically treated samples was expected.

**Figure 4.**
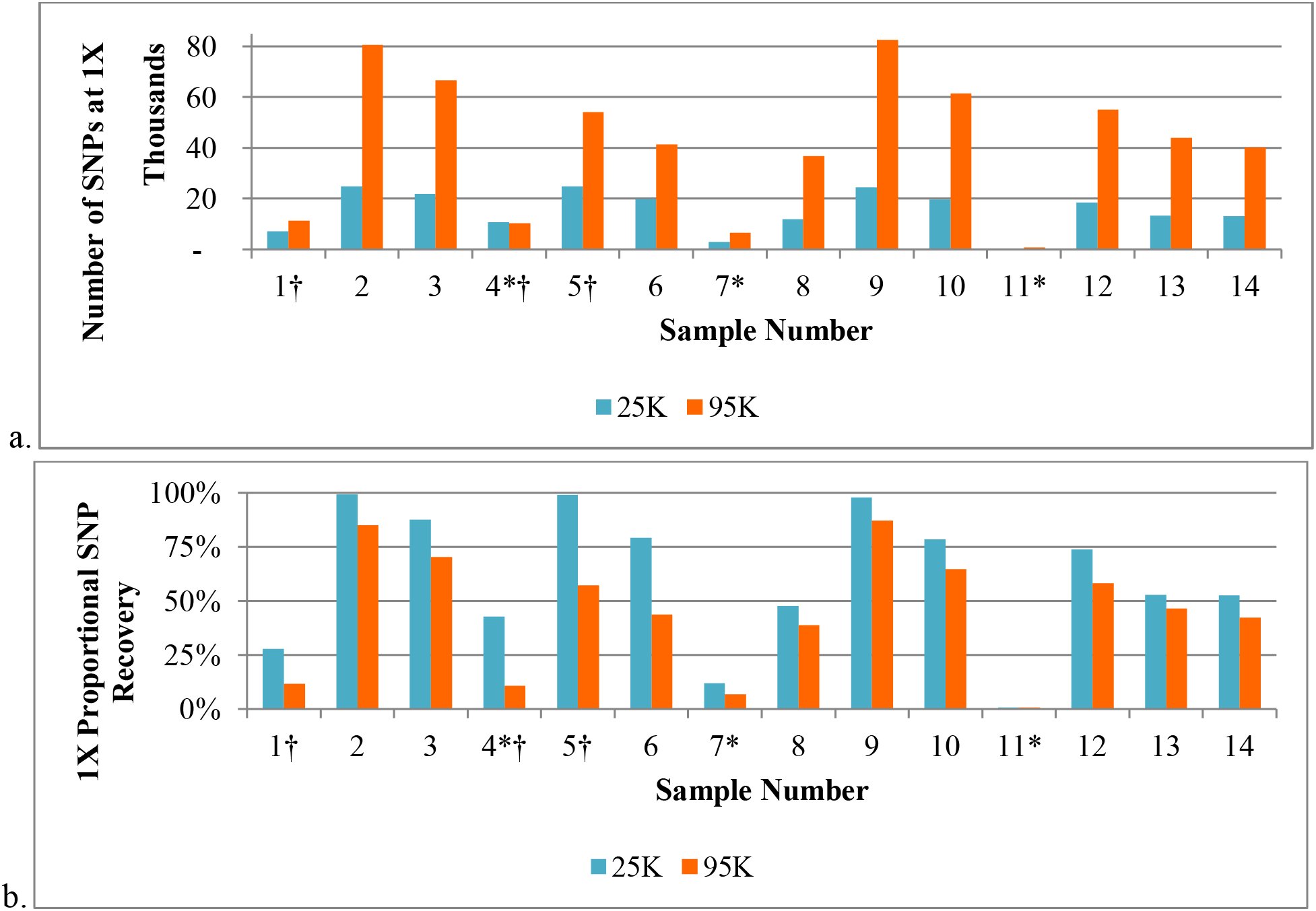
The (a) number and (b) proportion of single nucleotide polymorphisms (SNPs) recovered at ≥1X when 14 non-probative sample libraries were sequenced on the MiSeq. Samples indicated with an asterisk (*) are chemically treated (formalin preserved) specimens. The cross (†) indicates lower DNA input into the 95K capture than the 25K capture due to limited library volume.

When the coverage threshold was increased from 1X to 5X, the average number of SNPs decreased by more than 60% in the 25K dataset and by 75% in the 95K dataset (Figure 5). The usable data were further decreased by approximately 80-90% in both SNP panels to approximately 2,800 SNPs at a 10X threshold. Therefore, the proportion of targeted SNPs obtained was very small and insufficient for direct genotype calling using fixed coverage thresholds. Since the Parabon Fx software incorporates read depth into its probabilistic analytical approach, all ancestry and kinship predictions were performed using the 1X profiles.

**Figure 5.**
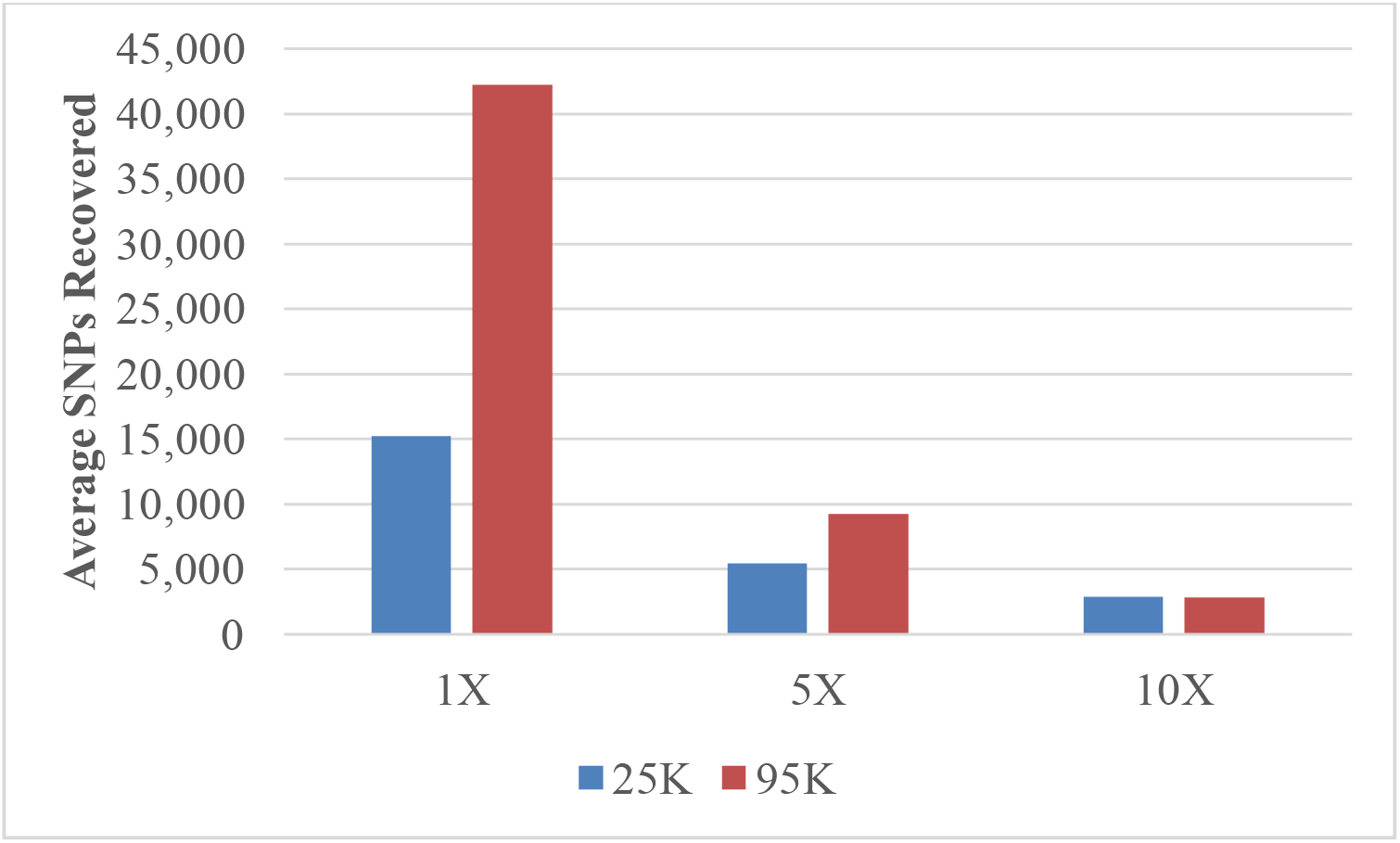
Average number of single nucleotide polymorphisms (SNPs) recovered for 14 skeletal samples, using coverage thresholds of 1X, 5X and 10X.

The two captured libraries sequenced on both the MiSeq and NextSeq (95K libraries for samples 2 and 5) yielded similar percentages of mapped reads on both instruments, yet smaller proportions were unique in the NextSeq data (Table S7; >49% for the MiSeq and <21% for the NextSeq). Regardless, the raw number of unique reads and SNP coverage per sample increased when the 95K capture product was sequenced on the NextSeq (Figure 6). This increase in SNP coverage was maintained in the NextSeq data when the minimum coverage threshold was increased to 5X or 10X. Higher coverage per SNP likely allows for improved kinship estimation. Therefore, sequencing on a higher throughput instrument may be beneficial due to the improved coverage, which is factored into the genotype likelihood. Although the two WGS libraries sequenced on the NextSeq generated more than 180,000 SNPs each at 1X coverage, this did not allow for accurate kinship prediction using the relatedness likelihood algorithm employed by the Parabon Fx software (discussed below). There were fewer than 30 SNPs each in the WGS data at 5X and 10X coverage thresholds; thus, multiple rounds of NextSeq sequencing would be required to improve SNP coverage.

**Figure 6.**
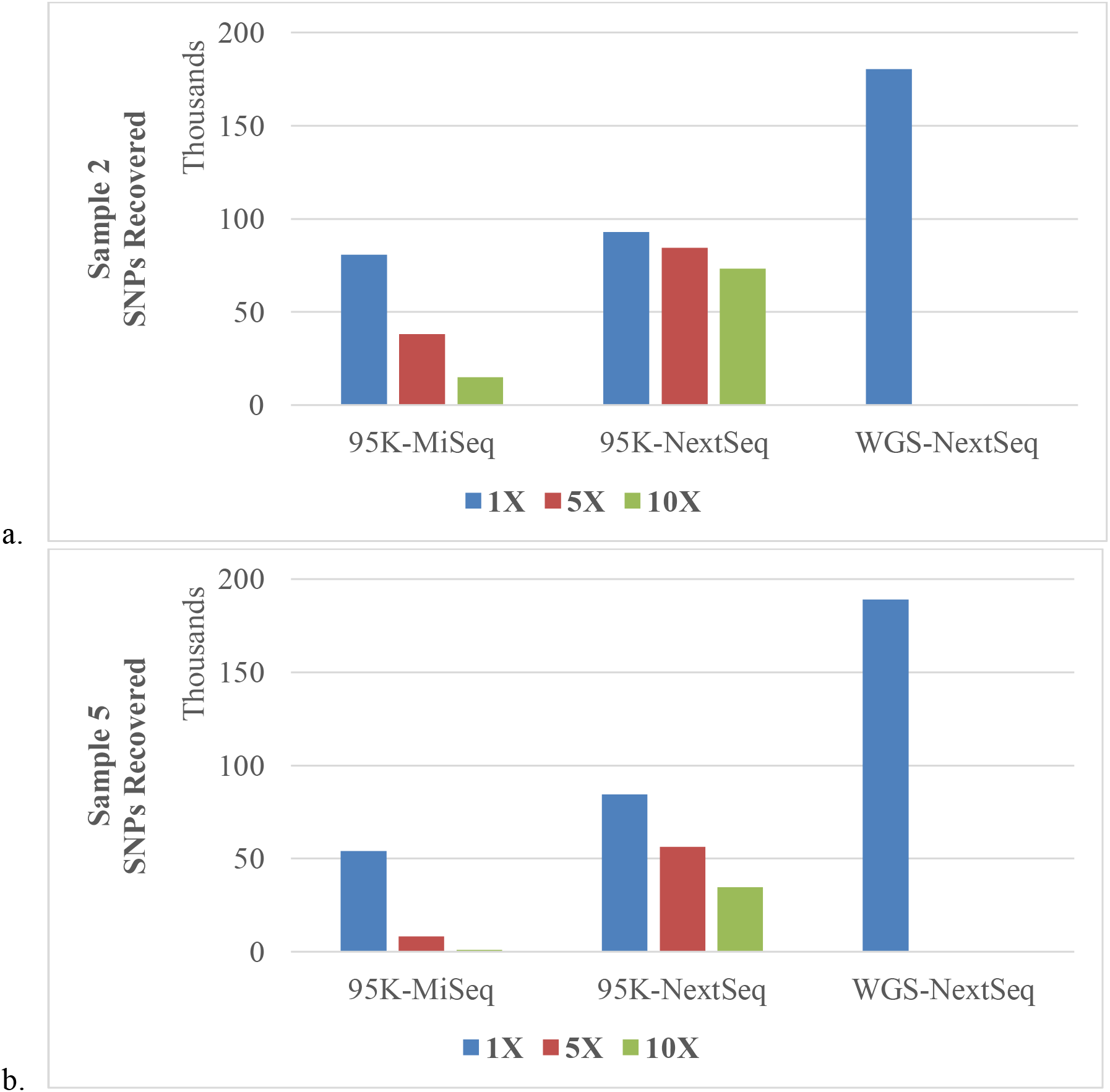
Number of single nucleotide polymorphisms (SNPs) recovered for samples 2 (a) and 5 (b) using 95K capture and whole genome sequencing (WGS), using coverage thresholds of 1X, 5X and 10X.

### 3.2 Microarray genotyping of family reference samples (FRS)

DNA quantities for FRS extracts were comparable across quantification methods (Table S8), averaging 1.75 ng/μl for Plexor HY and 1.70 ng/μl for Qubit HS. Overall, 35 of 39 (89.7%) FRS tested were successfully genotyped with microarray testing, yielding SNP call rates >96%. FRS 13-A demonstrated a moderate call rate (74%) but was still utilized in ancestry and kinship estimation. The three FRS with call rates <70% were not included in ancestry or kinship analysis. These four reference samples with call rates <96% yielded the lowest values using the Plexor HY DNA Quantification Kit (<210 pg/μl), indicating human DNA quantities below the recommended input for genotyping arrays. While not performed for this study, it is feasible that FRS could undergo SNP capture following DNA fragmentation and library preparation in order to obtain reference data. This would allow laboratories to implement a single processing method for SNP genotyping (i.e. capture) for all sample types. Fx offers capture-to-capture kinship analysis in addition to the capture-to-microarray analysis used for this study.

### 3.3 Parabon Fx ancestry prediction

Since allele frequencies of the targeted SNPs can vary substantially between population groups, failure to account for population differences in allele frequencies can impact the accuracy of kinship comparisons [37]. Ancestry prediction was thus performed on the case samples and the FRS within the Fx software using iAdmix [38], leveraging all reads mapping to targeted SNPs. The 36 FRS with sufficient call rates were determined to have European ancestry (Figure S4), consistent with the self-reported race provided. The ancestry of the known references was also used as a basis for the expected ancestry of the non-probative case samples. For all WGS, 25K capture and 95K capture samples, the ancestry predicted by Fx was European (Figure S5), consistent with the expected ancestry. Expansion of the populations tested, to include individuals of mixed race/ancestry, is needed in future research and validation efforts.

### 3.4 Parabon Fx kinship prediction

Kinship comparisons between the 1X non-probative case sample profiles and FRS are presented in Table 2 (21 related FRS) and Table S9 (15 unrelated FRS). The European allele frequency file was chosen for kinship estimation in all comparisons based on the European ancestry predictions obtained from both the FRS and the SNP capture data of the non-probative case samples. In the 25K SNP panel data, 17 of the 21 (81.0%) FRS were correctly predicted to be related to the corresponding case sample with LRs exceeding 10,000 (Table 2a). Precise kinship estimation, in which the correct relative *and* exact DOR were predicted, was possible for more than 70% of the 25K capture samples (15 of 21). Two samples were predicted to be related to the correct FRS but with an incorrect DOR. The other four of 21 FRS were correctly predicted to be related to the corresponding case sample from the 25K results, but the LR produced was below the 10,000 LR threshold utilized in this study to demonstrate strong statistical support for relatedness. The 95K data accurately associated the unknown sample with the correct FRS in 19 of 21 (90.5%) comparisons (Table 2b). The precise kinship prediction rate from the 95K data, in which the related FRS and DOR were correctly predicted, was then 81.0% (17 of 21) of pairwise comparisons. As also observed in the 25K data, the 95K kinship prediction identified the correct FRS but the wrong DOR for sample 2 / FRS 2-B and sample 13 / FRS 13-A (discussed below). The increased SNP recovery of the 95K panel permitted improved kinship inferences over the 25K panel, producing strong LR values for two of the four pairwise comparisons in which the 25K samples yielded LRs below the 10,000 LR threshold (sample 1 / FRS 1-B and sample 11 / FRS 11-A). Of note, the software was able to identify the correct FRS (11-A) for sample 11, a chemically treated sample, even though only 625 SNPs were recovered. FRS 11-A was correctly predicted to be a 1^st^ degree relative of sample 11, though the wrong relationship was indicated to be more likely (parent-offspring instead of full sibling). An increase in the SNP recovery for sample 11 through further rounds of sequencing would likely refine the kinship prediction. The two comparisons that did not produce kinship predictions exceeding the 10,000 LR threshold even with the 95K data were sample 1 / FRS 1-A and sample 7 / FRS 7-A. The poor SNP recovery (~11,000 and 6,500 SNPs for samples 1 and 7, respectively (Figure 4)), coupled with FRS of the furthest DOR tested (4^th^ degree), limited the strength of the kinship predictions for these two samples. Together the 25K and 95K capture data produced zero false positive relationships with LRs >10,000 in all 1,008 pairwise comparisons tested (504 per condition, excluding RBs), which includes pairwise comparisons with FRS that were unrelated to all case samples (Table S9). The NextSeq 95K data produced the same predicted relationships as the MiSeq 95K data, yet with higher LRs (sample 2 / FRS 2A: 1.83E+294; sample 2 / FRS 2B: 1.71E+118; sample 5 / FRS 5A: 1E+1084; sample 5 / FRS 5B: 1E+857).

**Table 2.** Likelihood ratios (LRs) were calculated for the highest predicted degree of relatedness (DOR) between each unknown and 21 family reference samples (FRS) compared to the two samples being unrelated. LRs greater than 1E+308 were reported as 1E(log LR) due to computational limitations. FRS 6-A, a relative of sample 6, failed genotyping and was excluded from kinship analysis. The remaining 15 unrelated FRS (excluding two failed FRS, 20-A and 21-A) were compared with the unknown samples and are presented in Table S9. Analysis was performed with data generated from SNP capture using 1X coverage data from the 25K (a) and 95K (b) panels when sequenced on the MiSeq, as well as NextSeq whole genome sequencing (WGS) for samples 2 and 5 (c). Samples are organized horizontally by DOR from closest to furthest (up to 4th degree). LRs above the 10,000 threshold are bolded; true relatives to each sample are designated by a thick border. Color indicates the degree of relatedness with the highest likelihood. In the tables, 1^st^ degree relatives are separated into parent/offspring (P/O) and full siblings (FS) as these precise relationships can be resolved within the software. Samples indicated with an asterisk (*) are chemically treated (formalin preserved) specimens. The cross (†) in the 95K panel (b) indicates lower DNA input into the 95K capture than the 25K capture due to limited library volume. P/O = yellow; FS = orange; 2^nd^ degree = green; 3^rd^ degree = blue; 4^th^ degree = pink; unrelated = white

| 25K |  | P/O |  | FS |  | 2nd |  |  |  |  | 3rd |  |  |  | 4th |  |
| --- | --- | --- | --- | --- | --- | --- | --- | --- | --- | --- | --- | --- | --- | --- | --- | --- |
|  |  | Sample Number |  |  |  |  |  |  |  |  |  |  |  |  |  |  |
|  |  | 4* | 14 | 11* | 12 | 3 | 5 | 6 | 13 | 14 | 2 | 8 | 9 | 10 | 1 | 7* |
| F<br>R<br>S<br># | 4-A | 4.55E+264 | 1 | 2 | 1 | 1 | 1 | 1 | 1 | 1 | 1 | 1 | 1 | 1 | 6 | 2 |
|  | 14-A | 1 | 1E+464 | 1 | 1 | 1 | 1 | 1 | 87 | 1E+464 | 1 | 1 | 1 | 1 | 1 | 1 |
|  | 11-A | 1 | 1 | 985 | 1 | 1 | 1 | 1 | 1 | 1 | 1 | 1 | 1 | 1 | 1 | 1 |
|  | 12-A | 1 | 1 | 1 | 1E+726 | 1 | 1 | 1 | 1 | 1 | 1 | 1 | 1 | 1 | 1 | 1 |
|  | 3-A | 1 | 1 | 1 | 1 | 1.66E+260 | 1 | 1 | 1 | 1 | 1 | 1 | 1 | 1 | 1 | 1 |
|  | 3-B | 1 | 1 | 1 | 1 | 3.65E+248 | 1 | 1 | 1 | 1 | 1 | 1 | 1 | 1 | 1 | 1 |
|  | 3-C | 1 | 1 | 1 | 1 | 3.07E+280 | 1 | 1 | 1 | 1 | 1 | 1 | 1 | 1 | 1 | 39 |
|  | 3-D | 1 | 1 | 1 | 1 | 8.14E+287 | 1 | 1 | 1 | 1 | 1 | 1 | 1 | 1 | 1 | 1 |
|  | 5-A | 1 | 1 | 1 | 1 | 1 | 1E+313 | 1 | 2 | 1 | 1 | 1 | 1 | 1 | 32 | 1 |
|  | 5-B | 1 | 1 | 1 | 1 | 1 | 1E+428 | 1 | 1 | 1 | 1 | 1 | 1 | 1 | 1 | 1 |
|  | 6-B | 1 | 1 | 1 | 1 | 1 | 1 | 7.64E+180 | 1 | 1 | 1 | 1 | 1 | 1 | 1 | 1 |
|  | 13-A | 1 | 1 | 6 | 1 | 1 | 1 | 1 | 5.37E+42 | 1 | 1 | 1 | 1 | 1 | 1 | 1 |
|  | 14-B | 1 | 1.70E+114 | 1 | 1 | 1 | 1 | 1 | 1 | 1.70E+114 | 1 | 1 | 1 | 1 | 1 | 2266 |
|  | 2-A | 1 | 1 | 2 | 1 | 1 | 1 | 1 | 1 | 1 | 1.09E+101 | 1 | 1 | 1 | 4 | 1 |
|  | 2-B | 1 | 1 | 1 | 1 | 1 | 1 | 1 | 1 | 1 | 1.44E+36 | 1 | 1 | 1 | 1 | 3 |
|  | 8-A | 1 | 1 | 1 | 1 | 1 | 1 | 1 | 1 | 1 | 1 | 1.06E+23 | 1 | 1 | 1 | 1 |
|  | 9-A | 1 | 1 | 1 | 1 | 1 | 1 | 10 | 1 | 1 | 1 | 1 | 6.08E+97 | 1 | 1 | 1 |
|  | 10-A | 1 | 1 | 1 | 1 | 1 | 1 | 1 | 1 | 1 | 1 | 1 | 1 | 1.73E+65 | 1 | 1 |
|  | 1-A | 1 | 1 | 1 | 1 | 1 | 1 | 1 | 1 | 1 | 1 | 1 | 1 | 1 | 255 | 1 |
|  | 1-B | 1 | 1 | 1 | 1 | 1 | 1 | 1 | 1 | 1 | 1 | 1 | 1 | 1 | 694 | 1 |
|  | 7-A | 1 | 1 | 1 | 1 | 1 | 1 | 1 | 1 | 1 | 1 | 1 | 1 | 1 | 1 | 3532 |

a.
| 95K |  | P/O |  | FS |  | 2nd |  |  |  |  | 3rd |  |  |  | 4th |  |
| --- | --- | --- | --- | --- | --- | --- | --- | --- | --- | --- | --- | --- | --- | --- | --- | --- |
|  |  | Sample Number |  |  |  |  |  |  |  |  |  |  |  |  |  |  |
|  |  | 4*† | 14 | 11* | 12 | 3 | 5† | 6 | 13 | 14 | 2 | 8 | 9 | 10 | 1† | 7* |
| F<br>R<br>S<br># | 4-A | 1.87E+219 | 1 | 1 | 1 | 1 | 1 | 1 | 1 | 1 | 1 | 1 | 1 | 1 | 1 | 1 |
|  | 14-A | 1 | 1E+1077 | 1 | 1 | 1 | 1 | 1 | 1 | 1E+1077 | 1 | 1 | 1 | 1 | 1 | 1 |
|  | 11-A | 1 | 1 | 1.04E+11 | 1 | 1 | 1 | 1 | 1 | 1 | 1 | 1 | 1 | 1 | 1 | 1 |
|  | 12-A | 4 | 1 | 1 | 1E+1723 | 1 | 1 | 1 | 1 | 1 | 1 | 1 | 1 | 1 | 1 | 1 |
|  | 3-A | 1 | 1 | 1 | 1 | 1E+594 | 1 | 1 | 1 | 1 | 1 | 1 | 1 | 1 | 1 | 2 |
|  | 3-B | 1 | 1 | 1 | 1 | 1E+617 | 1 | 1 | 1 | 1 | 1 | 1 | 1 | 1 | 3 | 1 |
|  | 3-C | 1 | 1 | 1 | 1 | 1E+598 | 1 | 1 | 1 | 1 | 1 | 1 | 1 | 1 | 1 | 1 |
|  | 3-D | 1 | 1 | 1 | 1 | 1E+700 | 1 | 1 | 1 | 1 | 1 | 1 | 1 | 1 | 1 | 1 |
|  | 5-A | 5 | 1 | 1 | 1 | 1 | 1E+434 | 1 | 1 | 1 | 1 | 1 | 1 | 1 | 1 | 1 |
|  | 5-B | 1 | 1 | 2 | 1 | 1 | 1E+520 | 1 | 1 | 1 | 1 | 1 | 1 | 1 | 1 | 1 |
|  | 6-B | 1 | 1 | 1 | 1 | 1 | 1 | 9.47E+241 | 1 | 1 | 1 | 1 | 1 | 1 | 1 | 1 |
|  | 13-A | 32 | 1 | 1 | 1 | 1 | 1 | 1 | 1.08E+116 | 1 | 1 | 1 | 1 | 1 | 1 | 11 |
|  | 14-B | 1 | 4.02E+305 | 1 | 1 | 1 | 1 | 1 | 1 | 4.02E+305 | 1 | 1 | 1 | 1 | 1 | 1 |
|  | 2-A | 1 | 1 | 1 | 1 | 1 | 1 | 1 | 1 | 1 | 1.12E+224 | 1 | 1 | 1 | 1 | 1 |
|  | 2-B | 1 | 1 | 1 | 1 | 1 | 1 | 1 | 1 | 1 | 2.59E+87 | 1 | 1 | 1 | 1 | 2 |
|  | 8-A | 1 | 1 | 3 | 1 | 1 | 1 | 1 | 1 | 1 | 1 | 1.17E+64 | 1 | 1 | 1 | 30 |
|  | 9-A | 1 | 1 | 1 | 1 | 1 | 1 | 1 | 1 | 1 | 1 | 1 | 4.05E+251 | 1 | 1 | 1 |
|  | 10-A | 1 | 1 | 1 | 1 | 1 | 1 | 1 | 1 | 1 | 1 | 1 | 1 | 2.77E+162 | 1 | 1 |
|  | 1-A | 1 | 1 | 3 | 1 | 1 | 1 | 1 | 1 | 1 | 1 | 1 | 1 | 1 | 208 | 1 |
|  | 1-B | 1 | 1 | 1 | 1 | 1 | 1 | 1 | 1 | 1 | 1 | 1 | 1 | 1 | 1.01E+04 | 1 |
|  | 7-A | 1 | 1 | 1 | 1 | 1 | 1 | 1 | 1 | 1 | 1 | 1 | 1 | 1 | 1 | 2340 |

b. C.
| WGS |  | 2nd | 3rd |
| --- | --- | --- | --- |
|  |  | Sample Number |  |
|  |  | 5 | 2 |
| F<br>R<br>S<br># | 4-A | 1 | 1 |
|  | 14-A | 1 | 1 |
|  | 11-A | 1 | 1 |
|  | 12-A | 1 | 1 |
|  | 3-A | 1 | 1 |
|  | 3-B | 1 | 1 |
|  | 3-C | 1 | 1 |
|  | 3-D | 1 | 1 |
|  | 5-A | 1E+424 | 1 |
|  | 5-B | 1E+495 | 1 |
|  | 6-B | 4.27E+05 | 1 |
|  | 13-A | 1 | 1 |
|  | 14-B | 1 | 1 |
|  | 2-A | 1 | 1.18E+170 |
|  | 2-B | 1 | 9.00E+114 |
|  | 8-A | 1 | 1 |
|  | 9-A | 1 | 1 |
|  | 10-A | 1 | 1 |
|  | 1-A | 1 | 1 |
|  | 1-B | 1 | 1 |
|  | 7-A | 1 | 1 |

The kinship results from the WGS data were not as robust as the capture data. In fact, WGS produced a false positive result between sample 5 and FRS 6-B, predicting a 4^th^ degree relationship with strong statistical support despite a lack of true relatedness between these individuals (Table 2c). This was the only false positive result observed in the WGS data (including the unrelated FRS in Table S9). Besides the false positive result, the four pairwise comparisons between samples 2 and 5 and their associated FRS predicted the correct FRS with strong statistical support but the wrong DOR, which was one additional degree removed from the expected relationship. It is important to note that the capture data (both 25K and 95K) also predicted one of these four relationships (sample 2 and FRS 2-B) to be one degree removed from the expected DOR and with strong statistical support. The weaker kinship results from WGS can most likely be attributed to the very low coverage data, which was primarily 1X (Figure 6). Since the read coverage of each allele is factored into the genotype likelihood, increased coverage as obtained in the 25K and 95K capture data may be important for precisely predicting the relationship and correct DOR. Improved read depth through even higher throughput sequencing of these WGS libraries may benefit the accuracy of the DOR predictions, although this would require production scale sequencing instrumentation (e.g., NovaSeq) that is not available in routine forensic laboratories. Hence, SNP capture may enable laboratories to perform distant kinship estimations from degraded bone samples using accessible resources, such as small benchtop sequencers.

As noted above, two of the kinship comparisons (sample 2 / FRS 2-B and sample 13 / FRS 13-A) predicted the correct FRS to be related, but the DOR was incorrect. This result was obtained for these two comparisons regardless of the SNP capture panel. In both cases, the predicted DOR was one degree more distant than the expected relationship. The LRs obtained for these incorrect relationships were exceptionally high (averaging 1.30 × 10^87^ for sample 2 / FRS 2-B, and 5.38 × 10^115^ for sample 13 / FRS 13-A). Additionally, samples 2 and 13 generated >24,000 and >13,000 SNPs, respectively. In the case of sample 2, FRS 2-B was expected to be a grand-niece (3^rd^ degree relative) of sample 2 but was predicted as a 4^th^ degree relative (e.g., great-grand-niece, half-grand-niece, or first cousin once removed). An incorrect classification of a 3^rd^ degree relative is not unexpected, as the transmission of genetic material is random, and there is variation in the proportion of DNA sharing within pairs of individuals of the same DOR [37]. Interestingly, sample 2 obtained high LRs (averaging 5.61 × 10^223^) for the correct DOR (3^rd^ degree) with its other FRS included in the study (FRS 2-A). Moreover, the relationship predicted between FRS 2-A and FRS 2-B was 3^rd^ degree, which is consistent with the reported relationship (first cousins) of the two FRS donors (Figure S6). Since the relationships between the sample 2 and FRS 2-A as well as the two FRS were predicted correctly, it is likely that the more distant DOR predicted for sample 2 and FRS 2-B may be the result of low allele sharing in the SNPs recovered rather than an issue with the genealogical record. By contrast, the reason for the incorrect DOR predicted for sample 13 was likely the reduced genotyping call rate of FRS 13-A (74%). It is therefore possible that a genotyping success threshold (e.g., >96%) should be established for FRS microarray data to permit reliable kinship estimation.

### 3.5 Analysis of control blanks

RBs were sequenced with the non-probative samples and analyzed to measure levels of background noise as well as check for possible contamination. Three of the four RB libraries were determined to be clean, with fewer than 100 SNPs covered at the 1X threshold (Table S7) and no SNPs were recovered from any of the RBs at the 5X or 10X thresholds. The 25K captured product of RB1 generated 1608 SNPs at 1X and required additional investigation. No SNPs were attributed to RB1 in the 95K data, indicating a lack of human DNA present and suggesting that the library was not contaminated. In fact, examination of the Bioanalyzer trace for RB1 showed a high adapter peak with no distinguishable capture product (Figure S7), providing additional support to the conclusion that RB1 was not contaminated during laboratory processing. Based on these findings, sequencing crosstalk is the most probable explanation for the reads attributed to RB1 in the 25K data [17,26,27].

To further investigate the likelihood of crosstalk, a capture-to-capture kinship comparison was performed between the RBs and the non-probative case samples (Table S10). The profile for RB1-25K matched sample 5 with a LR of 5.23 × 10^35^, indicating that the sequencing crosstalk likely came from this sample since sample 5-25K and RB1-25K were sequenced together in the same MiSeq run (Table S6). Crosstalk may have been caused by the high cluster density of this MiSeq run (Table S6) coupled with the sequence similarity between the indexes used for RB1 and sample 5 during library preparation (Table S11). A contaminated RB from suspected crosstalk could be resequenced with alternate samples to demonstrate the lack of contamination in the library. However, in routine practice implementation of an analytical threshold would assist in the classification of control cleanliness and/or sample data acceptance. In this particular instance of likely crosstalk, the proportional SNP recovery for RB1 was only 6.4% (1608 out of 24,999). Only one case sample (sample 11) dropped below this 1X proportional SNP recovery metric with <0.7% of SNPs covered in both the 25K data and 95K data (Table S7). Additional samples with low proportional SNP recoveries (<25%) correspondingly failed to produce accurate kinship results with strong statistical support using either capture panel (Table 2). Therefore, it may be worthwhile to evaluate proportional SNP recovery as an analytical threshold of low coverage data. For example, if less than 10% of the SNPs targeted were required for data analysis to be performed, sample 11 would be considered a failure and the data could not be utilized for kinship comparisons. Furthermore, RB1-25K data would not exceed this 10% proportional SNP recovery threshold and thus would be considered clean, effectively eliminating issues with low coverage background data introduced as a result of minimally multiplexed sequencing of captured samples and associated controls (e.g., RBs). Despite the proportional recovery in RB1 for the 25K sample set, none of the RBs matched to any of the FRS above the set LR>10,000 threshold under any of the conditions analyzed (Table S10).

## 4. Conclusions

This study demonstrates that adequate SNP data can be reliably obtained from aged skeletal samples by employing hybridization capture followed by NGS on a benchtop sequencer. The SNP recovery in combination with the Parabon Fx Forensic Analysis Platform were sufficient to permit accurate distant kinship predictions between severely degraded bone samples and known relatives. Though microarray testing will be successful for many forensic-type samples, DNA degradation and damage combined with low endogenous DNA content render aged skeletal remains recalcitrant to this traditional genotyping method. Also, as demonstrated in this study, reference samples may fail to generate reliable microarray data and thus require alternative genotyping methods due to lower DNA quality/quantity. Hybridization capture can be combined with a fragmenting library preparation method for high-quality samples, which would allow a single SNP genotyping method to be implemented for all sample types – lessening the validation burden and/or need for outsourcing of microarray testing for the laboratory.

Results from the capture data illustrate that, for most samples, the proportional SNP recovery is remarkably similar between the 25K and 95K panels. SNP recovery is thus proportional to the number of SNPs targeted in the panel. As it is impossible to determine which SNPs may be recovered from sample libraries, it benefits the kinship analysis to probe for the maximum number of SNPs possible. The Parabon Fx software facilitates this maximal SNP capture approach by implementing a probabilistic SNP profiling algorithm for low coverage sequence data that is tailored for degraded DNA samples with cytosine deamination. However, the maximum SNP probe capacity may be determined by factors such as the number of probes that can be effectively put in the capture assay, which must be balanced with cost and sequencing throughput. The present study demonstrates only a moderate (10%) increase in the proportion of precisely predicted relationships when increasing from the 25K to the 95K panel, which is roughly 50% more expensive. Although the cost of a custom capture panel is related to the bait design and number of reactions purchased, in the present study an order for 96 capture reactions would cost $17,760 for the 25K panel (four baits per SNP) versus $26,400 for the 95K panel (one to two baits per SNP). Laboratories pursuing hybridization capture should weigh the benefits of maximum SNP recovery with other factors including expected DOR, sample type, sequencing throughput, and budget.

A 25K SNP panel generated suitable data to accurately associate up to 3^rd^ degree relatives with the correct DOR prediction and strong statistical support. The recovery of additional SNPs with the 95K SNP panel permitted accurate association of up to 4^th^ degree relatives, the furthest DOR tested. WGS of two sample libraries sequenced on a NextSeq was less successful than SNP capture (MiSeq or NextSeq) of the same libraries. The WGS data produced one false positive result with strong statistical support, and it imprecisely predicted the DOR in four expected pairwise relationships. False positives, or adventitious hits, likely cannot be avoided, especially in large-scale databases [2] when dense marker sets are used [39]. In the capture data (both 25K and 95K panels), which produced higher coverage per SNP compared to WGS, no adventitious hits were generated with an LR threshold > 10,000 (the threshold utilized for this study). However, the capture data produced false negative results in four of 21 pairwise comparisons due to low LR values (<5,000). It is of larger consequence to misclassify two individuals as related than the alternative. Therefore it is imperative to choose laboratory methods and appropriate interpretation thresholds (e.g., LR, proportion of SNPs recovered) that minimize the number of false positives obtained.

In practice, it may be beneficial for the LR threshold to be based on simulations that determine the maximum expected value from unrelated individuals for a particular DOR. Additionally, the utility of calculating the posterior probability to provide statistical weight to the conclusions should be considered. Incorporation of the posterior probability into the interpretation of the results may be particularly useful in instances where the kinship analysis distinctly indicates relatedness, yet the predicted DOR is inconsistent with reported familial relationships. There are many explanations for noted differences between expected relationships and results obtained through genetic kinship analyses. The distribution of shared DNA varies widely among individuals of the same DOR, and the degree of overlap between different DORs increases as the relationships become more distant [40]. Alternatively, there may be issues with genealogical records, such as reporting flaws, non-paternity, or incorrect pedigrees. Policies should be in place for addressing these scenarios in light of sensitive family situations, while retaining the ability to identify the unknown individual (e.g., testing additional references) [2].

This approach of combining large-scale SNP capture with the Parabon Fx software tailored for degraded DNA analysis provided promising results for forensic genetics in this study, particularly for historical remains cases. SNP capture from aged, degraded skeletal samples may be especially impactful in cases where DNA degradation has prevented successful STR amplification and/or where there is a lack of paternal or maternal relatives necessary for the use of lineage markers. The ability to capture requisite SNP data overcomes STR limitations, expands the pool of eligible DNA sample donors suitable for kinship comparisons by enabling distant kinship predictions from living relatives, and removes the need to use lineage markers (mitochondrial DNA sequencing or Y-chromosomal STRs). Although further testing and validation studies are still required before implementation in a forensic laboratory, this study demonstrates the successful application of SNP capture combined with the Parabon Fx software to facilitate distant kinship estimation in decades-old unidentified remains cases.

## Disclaimer

The assertions herein are those of the authors and do not necessarily represent the official position of the United States Department of Defense, the Defense Health Agency, or its entities including the Armed Forces Medical Examiner System. Any mention of commercial products was done for scientific transparency and should not be viewed as an endorsement of the product or manufacturer.

## Acknowledgments

The authors would like to thank Jennifer Daniels-Higginbotham (SNA International, Armed Forces DNA Identification Laboratory) for laboratory analysis and assistance; Timmathy Cambridge (Armed Forces Medical Examiner System) for technical assistance; Amanda Sozer, (SNA International), Michael Fasano, Shairose Lalani, Lt Col Briones and COL Louis Finelli (Armed Forces Medical Examiner System) for administrative and logistical support.

